# A role for ATP Citrate Lyase in cell cycle regulation during myeloid differentiation

**DOI:** 10.1101/525303

**Authors:** Jess Rhee, Lauren A. Solomon, Rodney P. DeKoter

**Affiliations:** Department of Microbiology & Immunology and the Centre for Human Immunology, Schulich School of Medicine & Dentistry, Western University, London, Ontario, Canada N6A 5C1; Division of Genetics and Development, Children’s Health Research Institute, Lawson Research Institute, London, Ontario, Canada N6C 2R5

## Abstract

Differentiation of myeloid progenitor cells into macrophages is accompanied by increased PU.1 concentration and increasing cell cycle length, culminating in cell cycle arrest. Induction of PU.1 expression in a cultured myeloid cell line expressing low PU.1 concentration results in decreased levels of mRNA encoding ATP-Citrate Lyase (ACL) and cell cycle arrest. ACL is an essential enzyme for generating acetyl-CoA, a key metabolite for the first step in fatty acid synthesis as well as for histone acetylation. We hypothesized that ACL may play a role in cell cycle regulation in the myeloid lineage. In this study, we found that acetyl-CoA or acetate supplementation was sufficient to rescue cell cycle progression in cultured BN cells treated with an ACL inhibitor or induced for PU.1 expression. Acetyl-CoA supplementation was also sufficient to rescue cell cycle progression in BN cells treated with a fatty acid synthase (FASN) inhibitor. We demonstrated that acetyl-CoA was utilized in both fatty acid synthesis and histone acetylation pathways to promote proliferation. Finally, we found that *Acly* mRNA transcript levels decrease during normal macrophage differentiation from bone marrow precursors. Our results suggest that regulation of ACL activity is a potentially important point of control for cell cycle regulation in the myeloid lineage.

## Introduction

Highly proliferating cells, including cancers, preferentially use glycolysis over oxidative phosphorylation because, although it is less efficient at generating ATP, glycolysis is rapid and provides key metabolites for several biosynthetic pathways including nucleotide, amino acid, and fatty acid synthesis (1). This preferential use of glycolysis by cancer cells is known as the Warburg effect (1, 2). Glycolysis results in the production of citrate that can be exported from mitochondria to serve as the substrate for synthesis of acetyl-CoA by the enzyme ATP Citrate Lyase (ACL) (3). Acetyl-CoA is the required substrate of fatty acid synthase (FASN) in the first step of *de novo* fatty acid synthesis. Although normal cells and cancer cells can utilize exogenous lipids, FASN-mediated fatty acid synthesis is required to sustain the needs of highly proliferative cells (4). Many cancer types display increased endogenous fatty acid biosynthesis, regardless of the levels of extracellular lipids available (5). Inhibiting FASN is effective in limiting the growth and proliferation of cancer cells (4, 6).

As a central metabolite, there are several anabolic and catabolic pathways that can lead to the production of acetyl-CoA. These pathways can be located in mitochondria or in the cytoplasm (7). Within the mitochondria, acetyl-CoA is generated in the matrix by the pyruvate dehydrogenase complex, β-oxidation of fatty acids, and the catabolic metabolism of branched amino acids (7). In the cytoplasm, ACL is the central enzyme for production of acetyl-CoA from citrate. Acetyl-CoA synthetase 2 (ACSS2) can also generate acetyl-CoA from the substrate acetate that can be produced in the cell or imported into the cell (8).

Acetyl-CoA links lipid metabolism and histone acetylation to proliferation by being the midpoint in these two processes, with ACL produced acetyl-CoA diverted into both the fatty acid biosynthesis and histone acetylation pathways (9, 10). In addition to lipid biosynthesis, histone acetylation is also important for proliferation (9, 10). ACL is found in both the nucleus and cytoplasm, and RNA-interference-mediated silencing of ACL significantly reduces global histone acetylation (9). ACL is known to be upregulated in many cancers, and inhibition of ACL inhibits cancer cell proliferation (11–15). Therefore, ACL has an essential position in cellular processes, particularly lipid biosynthesis and histone acetylation, both of which influence cell cycle progression and proliferation (3, 9, 16).

PU.1 is a member of the E26-transformation-specific (ETS)-family of transcription factors and is essential for myeloid development (17, 18). PU.1 is expressed in hematopoietic stem cells (HSCs) and is further upregulated during myeloid differentiation (19, 20). Inhibition of PU.1 function in HSCs blocks subsequent myeloid differentiation (19, 21). During macrophage differentiation, PU.1 protein accumulates, correlating with increasing cell cycle length (22). Conversely, multiple studies have shown that decreased expression of PU.1 leads to increased cell cycle progression and eventually acute myeloid leukemia (AML) in mice (23–25). Our laboratory showed that cell cycle progression is inhibited by induction of PU.1 expression in a cultured myeloid cell line expressing low PU.1 concentration (Spi1^BN/BN^) (25, 26). Induction of PU.1 in inducible cells (iBN cells) resulted in macrophage differentiation and induction of microRNAs that targeted acetyl-CoA metabolism including *Acly* encoding ACL (27). We found that chemical inhibition of ACL activity was sufficient to block cell cycle progression in cultured BN cells (27).

The goal of the current study was to explore the mechanism(s) by which cell cycle is regulated in cultured BN cells through control of ATP Citrate Lyase (ACL) activity. We found that acetyl-CoA or acetate supplementation was sufficient to rescue cell cycle progression in cultured BN cells treated with an ACL inhibitor or induced for PU.1 expression. Acetyl-coA supplementation was also sufficient to rescue cell cycle progression in BN cells treated with a FASN inhibitor. Through lipid and histone extraction, we demonstrated that acetyl-CoA was utilized in both fatty acid synthesis and histone acetylation pathways to promote proliferation. However, in ACL inhibited cells, there was an increase in the amount of acetyl-CoA incorporated into lipids, suggesting that lipid biosynthesis may be a crucial pathway to promote proliferation. Finally, we found that *Acly* mRNA transcript levels decrease during normal macrophage differentiation from bone marrow precursors. Our results suggest that regulation of ACL activity is a potentially important point of control for cell cycle regulation in the myeloid lineage.

## Methods

### Mice

C57BL/6 mice were purchased from Charles River Laboratories (Saint-Constant, QC, Canada). All experiments were performed on protocols approved by the Western University Council on Animal Care.

### Cell culture

The BN and iBN inducible cell lines were previously described (25, 27). BN cells were cultured in Iscove’s modified Dulbecco’s medium (IMDM) (Wisent, St-Bruno, QC), with 1 ng/ml GM-CSF (Peprotech, QC), and additionally supplemented with 10% fetal bovine serum (FBS) (Wisent, St-Bruno, QC), penicillin (100 U/ml)/streptomycin (100 μg/ml) (Mediatech, Manassas, VA), L-glutamine (2 mmol/L) (Mediatech), 2-mercaptoethanol (5 x 10^-5^ M) (Sigma-Aldrich, St. Louis, MO). PU.1 induction experiments were performed by culture of iBN cells in 1.0 ng/ml GM-CSF in the presence of absence of 1000 ng/ml doxycycline for 48 hours. Additionally, BN/iBN cells were cultured in the presence or absence of 100 μM of acetyl-CoA (Sigma Aldrich, Oakville, ON), 25 mM of acetate (Sigma Aldrich, Oakville, ON), 55μM BMS303141 (Cedar Lane, Burlington, ON), or 10 μg/ml of C75 (Sigma Aldrich, Oakville, ON). Platinum-E (Plat-E) retroviral packaging cells were cultured in Dulbecco’s modified Eagle’s medium (DMEM; Wisent) supplemented with 10% fetal bovine serum (FBS) (Wisent), penicillin (100 U/ml)/streptomycin (100 μg/ml) (Mediatech), and L-glutamine (2mmol/L) (Mediatech).

### Cell Cycle Analysis

Cell cycle was analyzed by flow cytometry with an allophycocyanin (APC) BrdU Flow Kit according to the manufacturer’s protocol (BD Biosciences, Mississauga, ON). Cells were labeled with bromodeoxyuridine (BrdU) for 2 hours at 37°C. Cells were then incubated with the APC-conjugated anti-BrdU antibody using a 1:100 dilution. Cells were additionally stained with 7- amino-actinomycin D (7-AAD; BD Pharmingen) to determine cell cycle position. To stain cells with 7-AAD, cells were suspended in Dulbecco’s phosphate buffered saline (DPBS) supplemented with 5 mM ethylenediaminetetraacedic acid (EDTA) and 0.5% bovine serum albumin (BSA) and then incubated with 7-AAD.

### Flow Cytometry

Flow cytometry analysis was performed on single-cell suspensions of cells washed in flow cytometry buffer consisting of DPBS supplemented with 0.5 mM EDTA and 0.5% BSA. Antibodies directly conjugated to phycoerythrin (PE) against CD11b and C-kit were utilized to determine lineage-negative characteristics of extracted bone marrow cells. APC-BrdU was utilized for cell cycle analysis. Flow cytometric analysis and sorting was performed using a FACSCanto and FACSAria III, respectively (BD Immunocytometry Systems, San Jose, CA) at the London Regional Flow Cytometry Facility. Data was analyzed using FlowJo, version 10 (Tree Star, Ashland, OR).

### Retrovirus Production

Plat-E retroviral packaging cells (28) were used to generate retroviral supernatants using PEIPro transfection reagent (Polyplus, Ullkirch, France) at a 3:1 PEIPro/DNA ratio. Supernatant containing virus was collected 48 hours after transfection. Cells were infected by spinoculation by centrifugation at 3000 rpm for 3 hours at 32°C with polybrene at a final concentration of 8 μg/ml. After centrifugation, cells were washed and cultured for 48 hours to allow for retroviral integration and gene expression. Infection frequency was determined by flow cytometric analysis of green fluorescent protein (GFP).

### Bone Marrow Cell Isolation and Culture

Bone marrow cells were extracted from the femurs and tibias of C57Bl/6 mice. Bone marrow cells were washed three times with DPBS supplemented with 0.5 mM EDTA and 0.5% BSA. Bone marrow cells were stained using 1:100 dilutions of biotinylated antibodies recognizing CD11b, GR1, B220, and TER119 antibodies (BD Pharmingen, Mississauga, ON). The cells were then labelled with magnetic streptavidin microbeads according to the manufacturer’s instructions (Miltenyi Biotec, Auburn, CA). After washing, cells were lineage depleted using. LD columns (Miltenyi Biotec, Auburn, CA). Unfractionated and lineage-depleted (Lin^-^) cells were labelled with phycoerythrin (PE) conjugated antibodies for CD11b and c-Kit (BD Biosciences, Mississauga, ON), and analyzed by flow cytometry. Lin^-^ bone marrow cells were grown in culture with complete IMDM medium supplemented with 10 ng/ml M-CSF (Peprotech, QC).

### Tritium Culture and Scintillation Counting

BN cells were cultured with 0.5 μCi of [^3^H]-acetyl-CoA (Perkin Elmer, Waltham, MA) in 1 mL of complete media for 24 hours. Media was collected in separate tubes and the pellets were washed three times with PBS. The cell pellets were fully solubilized with 200 μl of a 3% solution of potassium hydroxide (KOH) (29). Solubilized cell pellets were then extracted for their lipid using 1ml of a 2:1 (v/v) chloroform:methanol solution, similar to the Folch method (without the neutralization step with acid/chloride salt) (Folch, Lees, & Stanley, 1957). Histones were prepared directly from cell pellets using a histone extraction kit according to the manufacturer’s instructions (Abcam, Toronto, ON). Disintegrations per minute (DPM) were determined using a LS6500 scintillation counter (Beckman Coulter, Ramsey, MN) using 10 μl from either solubilized cell pellets, lipid extracts and histone extracts.

### Reverse Transcriptase Quantitative PCR

RNA was isolated from cells using TRIzol Reagent (Life Technologies, Carlsbad, CA) and reverse-transcribed into cDNA using the iScript kit (Bio-Rad, Hercules, CA). Quantitative PCR (qPCR) was performed using iQ SYBR Green Supermix Kit (Bio-Rad) and a QuantStudio5 instrument (Applied Biosystems, Foster City, CA). Relative messenger RNA (mRNA) levels of *Acly* were normalized to *B2m* or *Adgre1* as reference genes and compared between samples using the comparative threshold cycle method (31). Primer sequences are listed in Supplemental Table 1.

### Statistical Analysis

Statistical significance was determined with ratio-paired *t* test or one-way analysis of variance (ANOVA) with Tukey’s multiple comparison using Prism 5 (GraphPad Software, La Jolla, CA, USA).

## Results

### Supplementation with acetyl-CoA or acetate rescues cell cycle arrest induced by inhibition of ATP Citrate Lyase in cultured myeloid cells

BN cells are myeloid precursor cells that are impaired for differentiation as a consequence of low PU.1 expression, and proliferate continuously in culture in response to granulocyte-macrophage colony-stimulating factor (GM-CSF) (26, 32). We previously showed that induction of PU.1 in inducible BN (iBN) cells resulted in downregulation of *Acly* encoding ATP Citrate Lyase (ACL), and that this regulation was likely indirectly mediated through induction of microRNAs (27). BMS303141 (BMS) is an effective inhibitor of ACL activity (33) and was sufficient to inhibit cell cycle progression in cultured BN cells (27). We set out to determine if supplementation of BN cells with acetyl-CoA could rescue BMS inhibition of cell cycle. BN cells were cultured with 55 μM BMS, a concentration found to result in half-maximal cell cycle inhibition as determined by the frequency of cells in S-phase (Fig. 1A, B, and data not shown). Addition of 100 μM acetyl-CoA to BMS-inhibited cultures resulted in a significant increase in the frequency of S-phase BN cells (Fig.1C, 1D). The sufficiency of acetyl-CoA to increase cell cycle progression of BN cells treated with BMS, suggests that this drug inhibits cell cycle progression by reducing the available cellular pool of acetyl-CoA.

**Figure 1.**
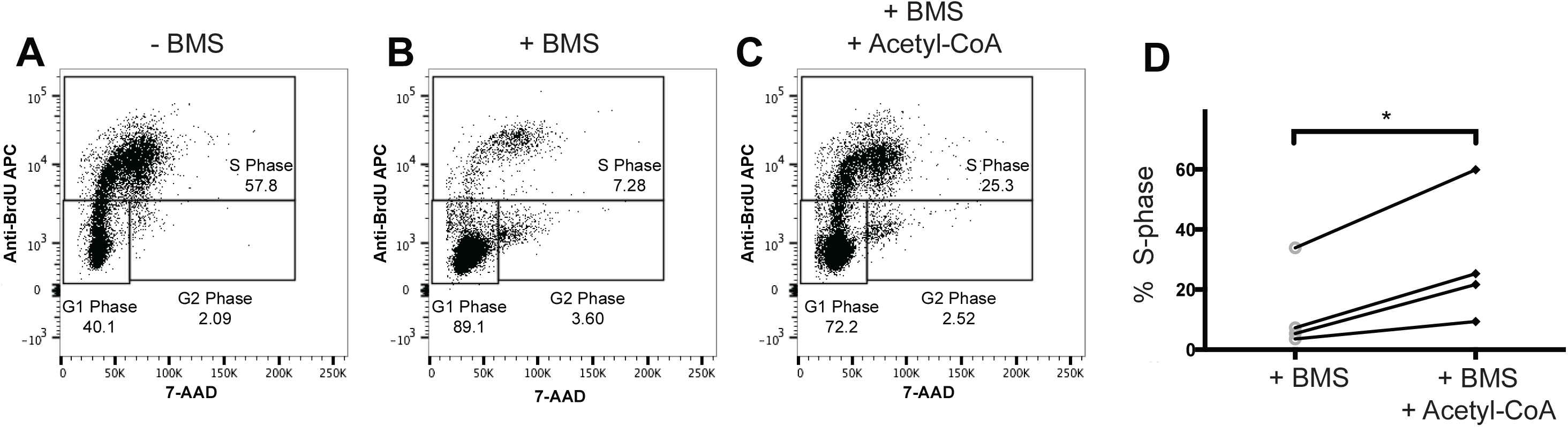
Regulation of cell cycle progression in cultured BN cells by ACL inhibition and acetyl-CoA supplementation. **A**) Representative cell cycle analysis of BN cells 48 hr after start of culture. **B**) Cell cycle analysis of BN cells cultured with 55μM BMS for 48 hr. **C**) Cell cycle analysis of BN cells cultured with 55μM BMS and 100 μM acetyl-CoA for 48 hr. **D**) Quantitation of the frequency of cells in S-phase for four experiments as shown in representative panels B and C. Statistical analysis was performed by paired t-test (* *p* < 0.05).

Next, acetate was used as a supplement since exogenous sources of acetate are able to be transported into the cell by monocarboxylate transporters (34). Acetyl-CoA synthetase 2 (ACSS2) can metabolize acetate into acetyl-CoA (8). Supplementation with 25 mM acetate significantly rescued proliferation of BN cells treated with 55 μM BMS (Fig. 2). This result suggests that BN cells could convert acetate to acetyl-CoA via the ACSS2 pathway to bypass the block induced by BMS treatment.

**Figure 2.**
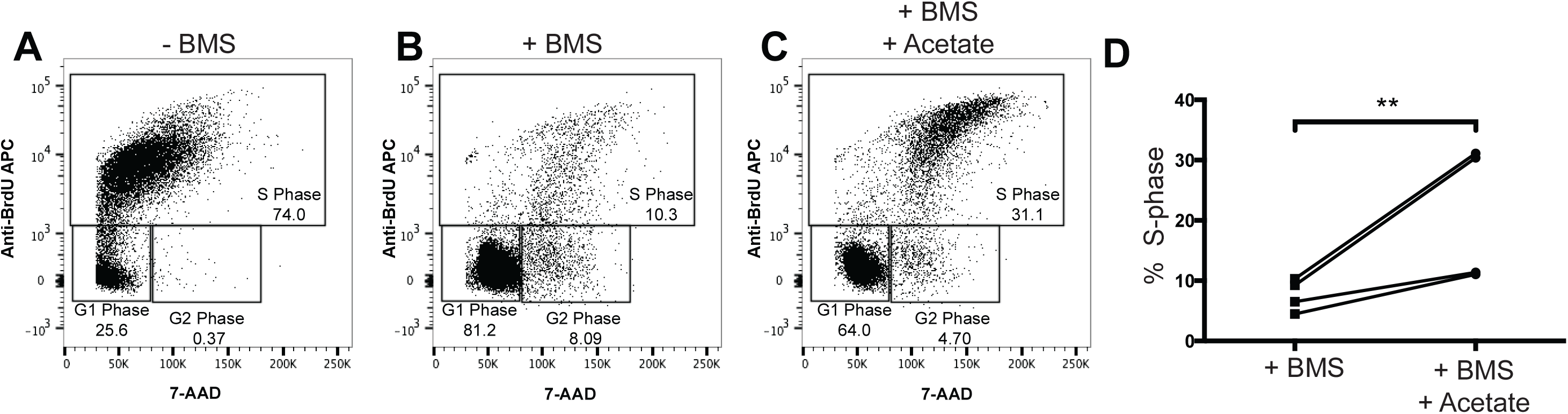
Regulation of cell cycle progression in cultured BN cells by ACL inhibition and acetate supplementation. **A**) Representative cell cycle analysis of BN cells 48 hr after start of culture. **B**) Cell cycle analysis of BN cells cultured with 55μM BMS for 48 hr. **C**) Cell cycle analysis of BN cells cultured with 55μM BMS and 25 mM acetate for 48 hr. **D**) Quantitation of the frequency of cells in S-phase for four experiments as shown in representative panels B and C. Statistical analysis was performed by paired t-test (*\*\* p* < 0.01).

### Supplementation with acetyl-CoA or acetate rescues cell cycle arrest induced by PU.1 induction in cultured myeloid cells

Our laboratory previously demonstrated that PU.1 induced iBN cells undergo cell cycle arrest accompanied by downregulation of *Acly* mRNA transcripts (27). Therefore, we wanted to determine if supplementing PU.1 induced iBN cells with acetyl-CoA or acetate was sufficient to rescue cell cycle arrest. Following induction of PU.1 in iBN cells with doxycycline, cell cycle progression was reduced (Fig. 3A, 2B). Supplementation with acetyl-CoA (Fig. 3C, 3D) and to a lesser extent acetate (Fig. 3E, 3F) significantly rescued PU.1-induced cell cycle arrest. These results suggest that PU.1 induces cell cycle arrest, at least in part, by down-regulation of production of acetyl-CoA.

**Figure 3.**
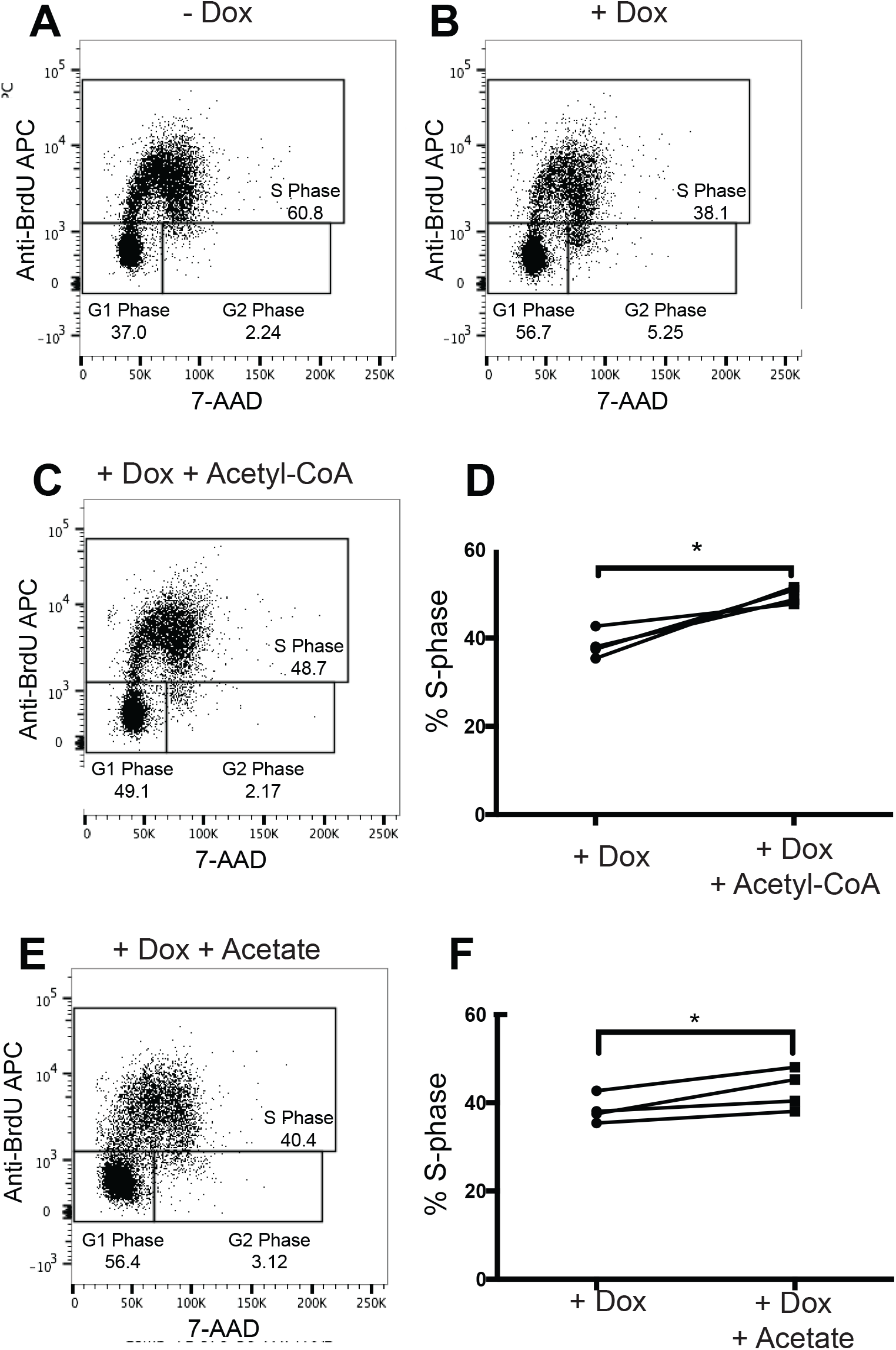
Regulation of cell cycle progression in cultured iBN cells by PU.1 induction and acetyl-CoA supplementation. **A**) Representative cell cycle analysis of iBN cells 48 hr after start of culture. **B**) Cell cycle analysis of iBN cells cultured with 1000ng/ml doxycycline for 48 hr. **C**) Cell cycle analysis of BN cells cultured with 1000ng/ml doxycycline and 100 μM acetyl-CoA for 48 hr. **D**) Quantitation of the frequency of cells in S-phase for four experiments as shown in representative panels B and C. Statistical analysis was performed by paired t-test (* *p* < 0.05). **E**) Cell cycle analysis of BN cells cultured with 1000ng/ml doxycycline and 25 mM acetate for 48 hr. **F**) Quantitation of the frequency of cells in S-phase for four experiments as shown in representative panels B and E. Statistical analysis was performed by paired t-test (* *p* < 0.05).

### Acetyl-CoA rescues cell cycle arrest from impaired FASNfunction

Given that acetate and acetyl-CoA supplementation were able to rescue cell cycle arrest from ACL inhibition via BMS and PU.1 induction via doxycycline, we next determined whether inhibition of fatty acid synthesis could arrest cell cycle in BN cells. C75 is an effective chemical inhibitor of FASN (35). Culture of BN cells with 10 μg/ml C75 efficiently inhibited cell cycle progression (Fig. 4A, 5B, 5D). Interestingly, supplementation of C75-treated BN cells with acetyl-CoA resulted in a significant rescue of cell cycle progression (Fig. 4C, 5D). This result suggests that inhibition of fatty acid synthesis is sufficient to impair cell cycle progression in cultured BN cells, and that supplementation with exogenous acetyl-CoA interferes with FASN inhibition by C75 to rescue fatty acid synthesis and cell cycle progression.

**Figure 4.**
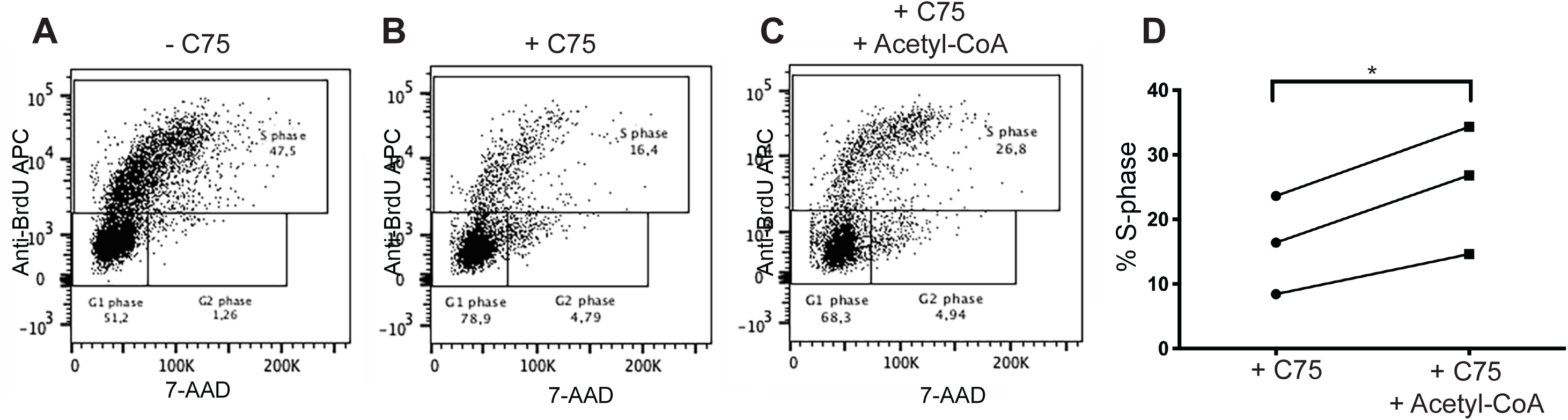
Regulation of cell cycle progression in cultured BN cells by FAS inhibition and acetyl-CoA supplementation. **A**) Representative cell cycle analysis of BN cells 48 hr after start of culture. **B**) Cell cycle analysis of BN cells cultured with 10 μg/ml C75 for 48 hr. **C**) Cell cycle analysis of BN cells cultured with 10 μg/ml C75 and 100 μM acetyl-CoA for 48 hr. **D**) Quantitation of the frequency of cells in S-phase for four experiments as shown in representative panels B and C. Statistical analysis was performed by paired t-test (* *p* < 0.05).

### Extracellular [^3^H]-Acetyl-CoA is incorporated into cells in a regulated manner

It has been shown that acetyl-CoA cannot enter cells by passive diffusion; and there are no descriptions of transporters that allow for import of extracellular acetyl-CoA (36). Therefore, we tested the import of extracellular acetyl-CoA into cells by the addition of extracellular [^3^H]-acetyl-CoA followed by liquid scintillation counting. BN cells were cultured with 0.5 μCi of [^3^H]-acetyl-CoA for 24 hours. After 24 hours, supernatants were collected and washed cell pellets were fully solubilized using 3% KOH followed by liquid scintillation counting (29). Solubilized cell pellets of iBN cells incubated with [^3^H]-acetyl-CoA incorporated a significant amount of ^3^H compared to the negative control (Fig. 5A). As expected, most [^3^H]-acetyl-CoA remained in the supernatant (Fig. 5A).

**Figure 5.**
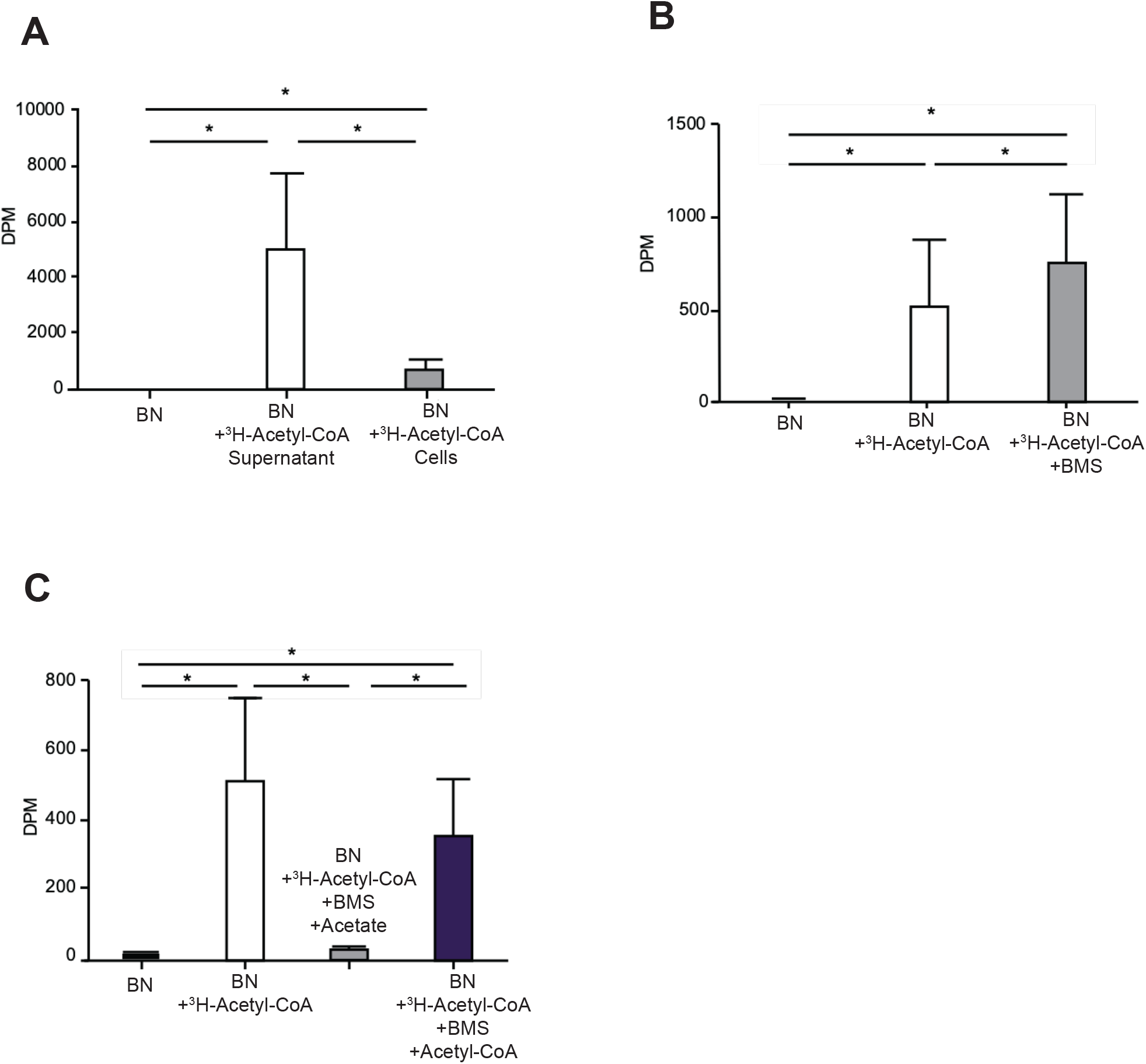
Regulated incorporation of [^3^H]-acetyl-CoA into cultured BN cells. **A**) Incorporation of [3H]-acetyl-CoA into cultured BN cells. BN cells were incubated for 24 hr with 0.5 μCi [^3^H]-acetyl-CoA. DPM were determined in the supernatant and solubilized cell pellets. Statistics were determined using one-way ANOVA with Tukey’s multiple comparisons test, *p* < 0.05 (n=6). **B**) Increased incorporation of [3H]-acetyl-CoA into cultured BN cells upon ACL inhibition. BN cells were treated with (right bar) or without (middle bar) 55 μM BMS and incubated for 24 hr with 0.5 μCi [^3^H]-acetyl-CoA. DPM were determined in solubilized cell pellets. Data is presented as mean ± SEM, one-way ANOVA with Tukey’s multiple comparisons test, *p* < 0.05, n=7. **C**) Decreased incorporation of [3H]-acetyl-CoA into cultured BN cells upon BMS and acetate but not BMS and Acetyl-CoA supplementation. BN cells were supplemented with either 25 mM acetate (third bar) or 100 μM acetyl-CoA (fourth bar) at the same time as 55 μM BMS and incubated for 24 hr with 0.5 μCi [^3^H]-acetyl-CoA. DPM were determined in solubilized cell pellets. Data are presented as mean ± SEM. Statistics were determine using one-way ANOVA with Tukey’s multiple comparisons test, * *p* < 0.05 (n=5 experiments).

To determine if uptake of [^3^H]-acetyl-CoA can be actively regulated, we determined if ACL inhibition affected [^3^H]-acetyl-CoA incorporation. BN cells were cultured in media with 55 μM of BMS to inhibit ACL activity and 0.5 μCi of [^3^H]-acetyl-CoA. Interestingly, culture with BMS resulted in a significant increase in the incorporation of [^3^H]-acetyl-CoA in BN cells (Fig. 5B). This result suggests that the [^3^H]-acetyl-CoA import mechanism is actively modulated by intracellular acetyl-CoA concentrations.

Next, we set out to determine if supplementation with acetate could affect uptake of [^3^H]-acetyl-CoA in BN cells. Interestingly, supplementation with 25mM acetate dramatically reduced the amount of [^3^H]-acetyl-CoA incorporation by BMS-treated BN cells (Fig. 5C). In contrast, supplementation with 100 μM acetyl-CoA did not significantly reduce the amount of [^3^H]-acetyl-CoA incorporation by BMS-treated BN cells (Fig. 5C). These results suggest that acetate, but not exogenous acetyl-CoA, is sufficient to compensate for the need to import additional acetyl-CoA upon BMS inhibition. Taken together, these results suggest that BN cells actively take up and incorporate [^3^H]-acetyl-CoA from the exogenous environment of the cells, and that supplementation with acetate results in production of enough acetyl-CoA to prevent detection of this incorporation from the exogenous environment.

### [^3^H]-Acetyl-CoA is incorporated into both lipids and histones

Next, we set out to gain insight into the class of mechanism by which acetyl-CoA regulates cell cycle progression in BN cells. Acetyl-CoA might be able to regulate cell cycle progression by providing the biological substrates to sustain lipid biosynthesis and proliferation. A second non-mutually exclusive possibility is that acetyl-CoA concentration is a limiting factor for histone acetylation to promote gene activation of genes involved in cell cycle regulation (9). To determine which of these two pathways is involved in acetyl-CoA incorporation, BN cells were incubated for 24 hours with 0.5 μCi of [^3^H]-acetyl-CoA. Lipids were enriched using the Folch extraction method (30) and histones were enriched using a histone extraction kit before liquid scintillation counting. The results showed that [^3^H]-acetyl-CoA was incorporated into both lipids and histones, although quantitatively more was incorporated into lipids (Fig. 6A). This result suggests that both pathways may be active in BN cells, although lipid biosynthesis may be quantitatively more important.

**Figure 6.**
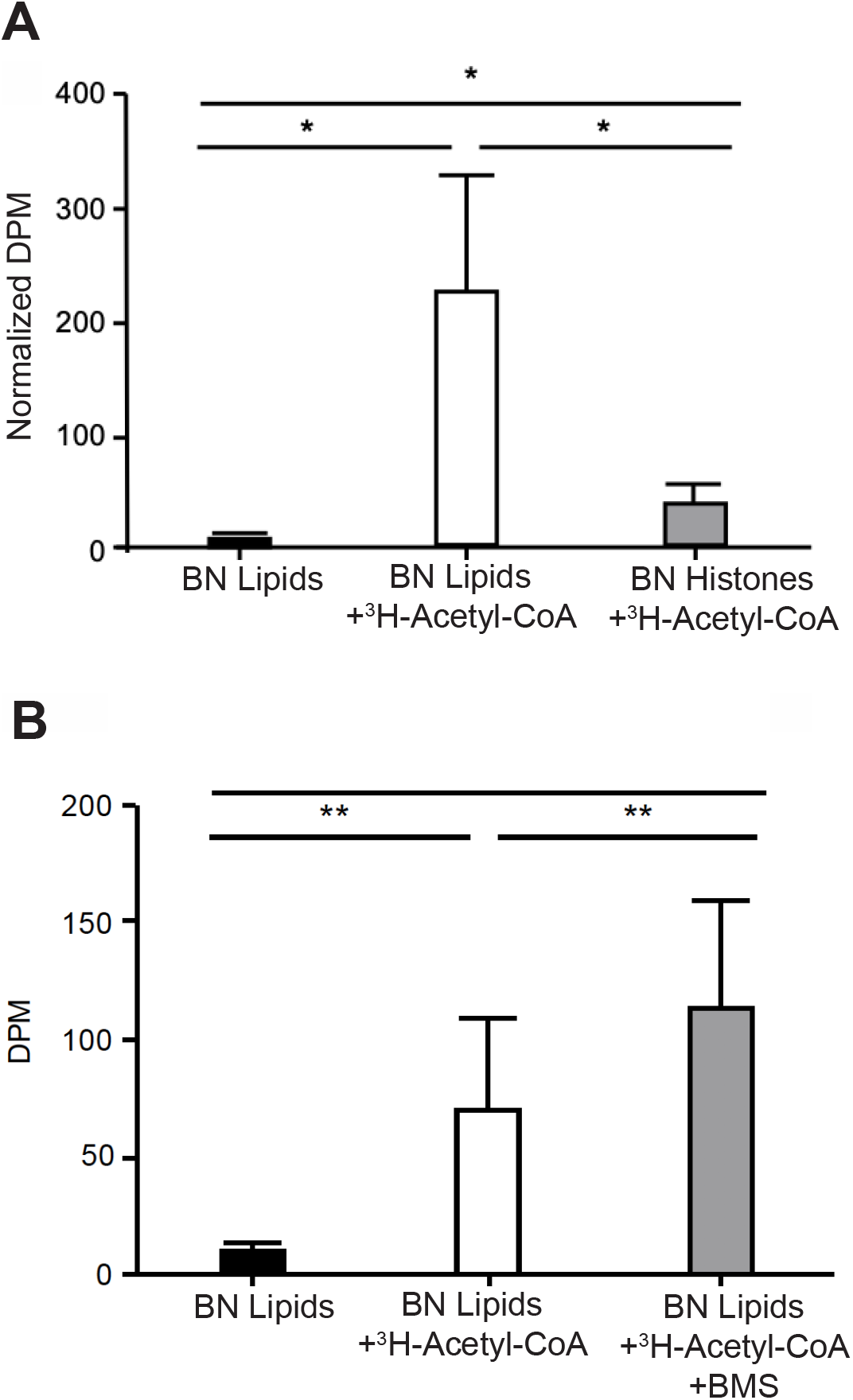
[^3^H]-acetyl-CoA is incorporated in both lipids and histones. **A**) BN cells were incubated with [^3^H]-acetyl-CoA for 24 hours followed by enrichment of lipids or histones as described in Materials & Methods. Normalized DPM was determined using liquid scintillation counting and correction for volume. Data are presented as mean ± SEM, one-way ANOVA with Tukey’s multiple comparisons test, *p* < 0.05, n=5. **B**) Increased incorporation of [^3^H]-acetyl-CoA into lipids upon ACL inhibition. BN cells were treated with (right bar) or without (middle bar) 55 μM BMS and incubated for 24 hr with 0.5 μCi [^3^H]-acetyl-CoA. DPM were determined in lipid extracts. Data are presented as mean ± SEM, one-way ANOVA with Tukey’s multiple comparisons test, * *p* < 0.05, ** *p* < 0.01, n=5 experiments.

In order to confirm and extend the result shown in Fig. 5B that ACL inhibition by BMS induces increased uptake of [^3^H]-acetyl-CoA, we repeated this experiment and additionally performed lipid extraction on BN cells treated with or without BMS. The results showed that BMS treatment increased the amount of [^3^H]-acetyl-CoA incorporation in lipids in BN cells compared to cells not treated with BMS (Fig. 6C). This result suggests that when acetyl-CoA concentrations are limiting, cells divert this molecule towards the pathway of lipid biosynthesis. Taken together, these results are consistent with the idea that ACL inhibition impairs cell cycle/proliferation through decreased lipid biosynthesis, and inhibition can be partially reversed by the exogenous addition of acetyl-CoA.

### Decrease in Acly mRNA transcript levels duringM-CSF-dependent macrophage differentiation

Finally, we wanted to determine whether *Acly* levels decrease during differentiation of myeloid progenitor cells into macrophages. If decreasing ACL levels are involved with decreasing cell cycle progression, then we would expect *Acly* mRNA transcript levels to decrease during macrophage differentiation. To test this idea, bone marrow cells from C57Bl/6 mice were prepared and lineage depleted (Lin^-^) using antibodies for CD11b, GR-1, B220, and TER119 to remove mature macrophages, granulocytes, B cells, and erythrocytes, respectively (Fig. 7A). CD11b+ myeloid cells were efficiently depleted in the Lin^-^ bone marrow cell fraction (Fig. 7B). Lin^-^ cells were cultured in 10 ng/ml M-CSF for 6 days. Adherent cells were detectable at day 2 and cells with macrophage morphology were visible by days 4 and 6 (Fig. 7C). RNA was prepared from adherent cells on days 2, 4, and 6. RT-qPCR analysis was performed to determine the steady-state mRNA transcript levels for *Acly* relative to either *B2m* (encoding β2 microglobulin) or *Adgre1* (encoding F4/80) as reference genes. There was a significant decrease in the level of *Acly* mRNA transcripts with reference to either *B2m* (Fig. 7D) or *Adgre1* (Fig. 7E). This observation suggests that mRNA transcript levels for *Acly* decrease during macrophage differentiation, consistent with a potential role in regulation of cell cycle progression in the myeloid lineage.

**Figure 7.**
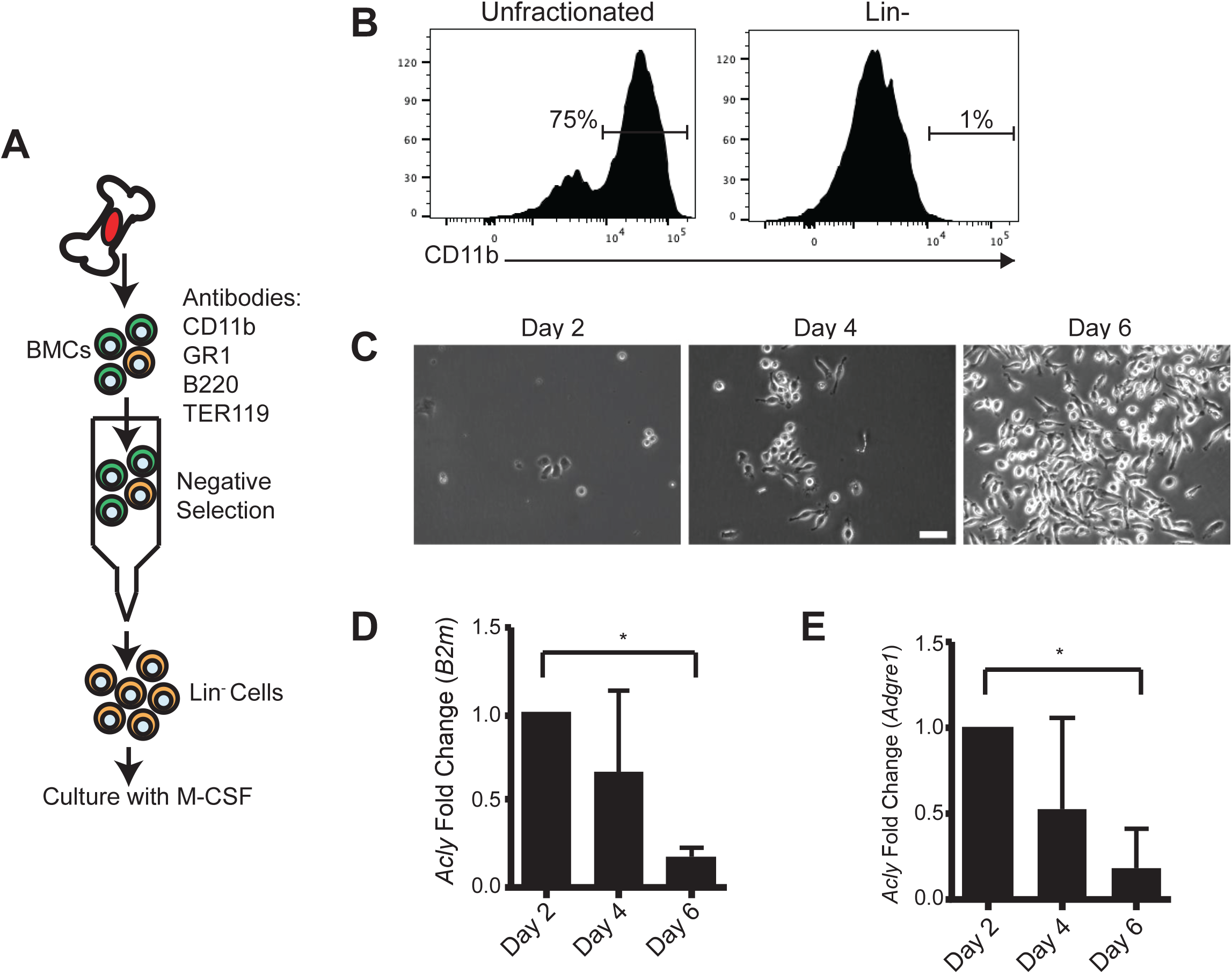
Decreased *Acly* mRNA transcript levels during macrophage differentiation. **A**) Schematic for enrichment of Lineage negative (Lin^-^) myeloid progenitor cells from bone marrow using magnetic bead depletion. **B**) Depletion of CD11b+ cells from bone marrow. Left panel shows frequency of CD11b+ cells in unfractionated bone marrow. Right panel shows frequency of CD11b+ cells in Lin^-^ bone marrow. **C**) Generation of macrophages by culture in complete media containing 10 ng/ml M-CSF. Representative photomicrographs were taken at 2 (left panel), 4 (middle panel), and 6 (right panel) days. Scale bar indicates 20 μm. **D, E**) *Acly* mRNA transcript levels decrease during macrophage differentiation. Using RT-qPCR, mRNA levels of *Acly* in the column separated BMCs were determined for day 2, 4, and 6 time points. *Acly* transcript levels were normalized to β2m (D) or *Adgre1* (E). Statistical analysis was performed using two-way ANOVA, * *p* < 0.05, n=3 experiments.

## Discussion

In this study, we showed that acetyl-CoA or acetate supplementation was sufficient to rescue cell cycle progression in cultured BN cells treated with BMS or induced for PU.1. Acetyl-CoA supplementation was sufficient to rescue cell cycle progression in cultured BN cells treated with C75. Through lipid and histone extraction, we were able to demonstrate that acetyl-CoA was utilized in both fatty acid synthesis and histone acetylation pathways to promote proliferation. In ACL inhibited cells there was an increase in the amount of acetyl-CoA incorporated into lipids, suggesting that lipid biosynthesis may be a crucial pathway to promote proliferation. Finally, we found that *Acly* mRNA transcript levels decrease during normal macrophage differentiation from bone marrow precursors. Our results suggest that downregulation of ACL activity is a potentially important point of control for cell cycle regulation in the myeloid lineage.

Acetate can be actively transported into cells and metabolized to acetyl-CoA by the enzyme acyl-CoA synthetase short-chain family member 2 (ACSS2) (8). However, there are no described mechanisms for active transport of acetyl-CoA (36). Acetyl-CoA transporters have been discovered in other subcellular compartments, notably acetyl-CoA transporters-1 (AT-1), which is expressed on the endoplasmic reticulum (ER) and mediated acetyl-CoA transport into the ER lumen (37–39). In our experiments we found that exogenous supplementation of cell culture media with acetyl-CoA could rescue cell cycle arrest by chemical inhibition of ACL or FASN. Furthermore, [^3^H] acetyl-CoA was incorporated into cellular lipids in a manner that was increased by inhibition of ACL. These results strongly suggest active transport of acetyl-CoA into cultured BN cells. Given that the AT-1 transporter is found to be localized on the ER, we speculate that AT-1 could be exported to the Golgi apparatus, and from the Golgi to the cell membrane (40). A second possibility is that acetyl-CoA enters BN cells through active phagocytosis or pinocytosis. BN cells express PU.1, although at low concentration, and the pattern of gene expression in these cells suggests that they are immature myelomonocytic cells that are capable of phagocytosis (25). An important question is whether either lipid biosynthesis or histone acetylation is the “sensor” to signal for cell cycle regulation by ACL inhibition. BMS treated iBN cells had higher [^3^H]-acetyl-CoA incorporation in their lipids than histones, suggesting that lipid biosynthesis may be the preferential pathway by which extracellular acetyl-CoA rescues iBN cells from cell cycle arrest. Consistent with lipid synthesis being the predominant mechanism by which ACL inhibition blocks cell cycle progression, one study utilized shRNA to silence *Acly* expression and showed reduced proliferation in these *Acly* silenced cells when grown in reduced lipid conditions (14). This study showed that supplementation with fatty acid, oleic acid, and acetate was sufficient to rescue cell cycle (14). Therefore, taken together, these results suggest that acetyl-CoA as a supplement is able to restore cell cycle progression in BMS treated BN cells by restoring the acetyl-CoA available in these cells for lipid biosynthesis to allow for cell cycle progression.

Previous studies suggested a link between PU.1 concentration and regulation of cell cycle progression. Small reductions in PU.1 concentration lead to increased proliferation and reduced differentiation of hematopoietic stem cells and myeloid progenitor cells (24, 26, 41). Re-introduction of PU.1 into proliferating cells expressing low PU.1 concentration rapidly induces cell cycle arrest and differentiation (23, 25). Induction of cell cycle arrest by PU.1 is accompanied by reduced *Acly* mRNA transcript levels (27). We showed in the current study that *Acly* mRNA transcript levels also decreased during normal M-CSF-dependent macrophage differentiation (Fig. 8). Therefore, we speculate that macrophage differentiation from myeloid progenitors involves several feedback loops involving PU.1 and *Acly.* Reduced *Acly* mRNA and ACL protein levels would lead to increased cell cycle length due to reduced lipid biosynthesis. Increased cell cycle length promotes PU.1 protein accumulation (22). High PU.1 concentration would then promote macrophage differentiation as marked by increased expression by genes such as *Adgre1* and expression of microRNAs targeting *Acly* (27). This would feed back to further decrease ACL levels to promote cell cycle arrest and macrophage differentiation.

In conclusion, it is well documented that nutrient depletion impairs cell cycle progression in normal or cancerous cells. However, there is still little known about the sensor(s) that transmit information about nutrient status to the cell cycle clock (42, 43). Our results suggest that regulation of the acetyl-CoA pool in cells may be an important mechanism to control cell cycle in developing myeloid cells. The size of the acetyl-CoA pool can be sensed by multiple mechanisms including through the rate of lipid biosynthesis or by histone acetylation (44). An important future direction will be to further explore the pathway by which the size of the intracellular pool of acetyl-CoA regulates the cell cycle clock in myeloid cells.

## Acknowledgements

We thank Kristin Chadwick and the London Regional Flow Cytometry Core Facility for assistance with flow cytometric analysis. We thank Drs. Murray Huff and Bryan Heit (Western University) for helpful advice and discussion. This work was supported by a Canada Graduate Scholarship-Masters to J.R. and by a Natural Sciences and Engineering Research Council Discovery Grant to R.D. (Grant 04749-2010).

---

**Supplemental Table 1.**
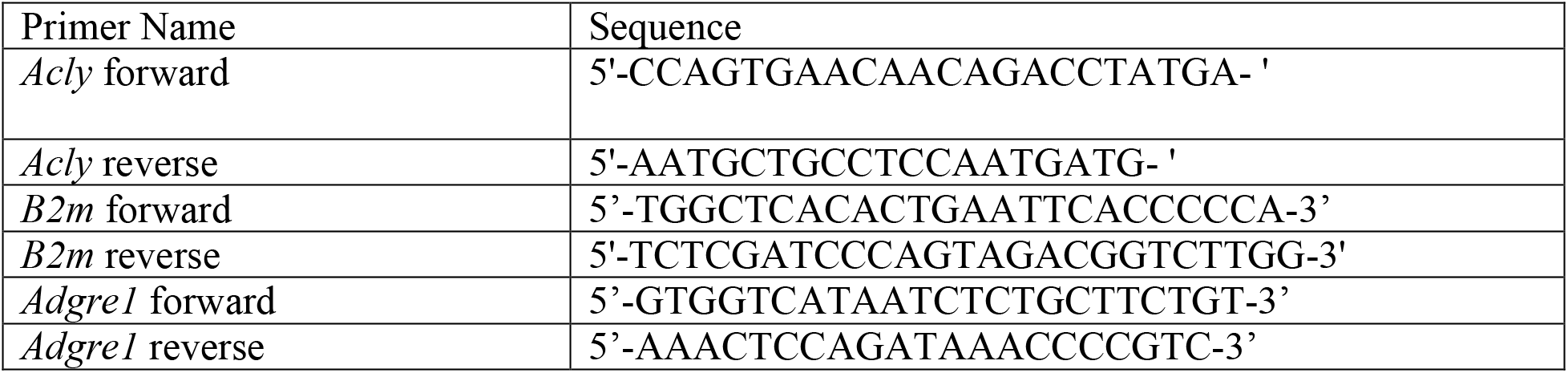
Primer sequences.

